# Characterization of the bHLH Family of Transcriptional Regulators in the ACOEL *S. roscoffensis* and their Putative Role in Neurogenesis

**DOI:** 10.1101/237388

**Authors:** E Perea-Atienza, S.G. Sprecher, P Martínez

## Abstract

**Background:** The basic Helix loop helix (bHLH) family of transcription factors is one of the largest superfamilies of regulatory transcription factors and are widely used in eukaryotic organisms. They play an essential role in a range of metabolic, physiological, and developmental processes, including the development of the nervous system (NS). These transcription factors have been studied in many metazoans, especially in vertebrates but also in early branching metazoan clades such as the cnidarians and sponges. However, currently very little is known about their expression in the most basally branching bilaterian group, the xenacoelomorphs. Recently, our laboratory has characterized the full complement of bHLH in the genome of two members of the Xenacoelomorpha, the xenoturbellid *Xenoturbella bocki* and the acoel *Symsagittifera roscoffensis*. Understanding the patterns of bHLH gene expression in members of this phylum (in space and time) provides critical new insights into the conserved roles of the bHLH and their putative specificities in this group. Our focus is on deciphering the specific roles that these genes have in the process of neurogenesis.

**Results:** Here, we analyze the developmental expression of the whole complement of bHLH genes identified in the acoel *S. roscoffensis.* Based on their expression patterns several members of bHLH class A appear to have specific conserved roles in neurogenesis, while other class A genes (as well as members of other classes) have likely taken on more generalized functions. All gene expression patterns are described in embryos and early juveniles.

**Conclusion:** Our results suggest that the main roles of the bHLH genes of *S. roscoffensis* are evolutionarily conserved, with a specific subset dedicated to patterning the nervous system: SrAscA, SrAscB, SrHes/Hey, SrNscl, SrSrebp, SrE12/E47 and SrOlig.

## INTRODUCTION

Xenacoelomorpha is a phylum constituted by small, mostly marine, benthic worms that share a relatively “simple” morphology (reviewed in [1, 2]). They are bilaterally symmetrical and their bodies are covered by a ciliated epithelium with a mouth being the only digestive opening to the exterior. Common features found in other metazoans such as the presence of circulatory or excretory systems, anus and coelom are completely absent in this group [3–5]. Xenacoelomorpha is divided into three clades: Xenoturbellida, Nemertodermatida and Acoela [6]. The latter two are grouped into the clade Acoelomorpha, the sister group of Xenoturbellida. For a long time, the phylogenetic affinities of these clades have been a matter of intense debate; see[6–9]. However, the latest phylogenetic analysis carried out by Canon and colleagues [10] seemed to resolve this conflict, proposing, based on analysis with strong support, a monophyletic Xenacoelomorpha as the sister group to all remaining bilaterians.

There are several reasons why xenacoelomorphs are an interesting set of biological systems in which to carry out comparative molecular and developmental studies. They belong to a monophyletic group (sharing a common ancestor) which members show a high diversity in the complexity of many regulatory families and in the organization of anatomical architectures (i.e. nervous system, arrangement of musculature, position of the mouth, the morphology of copulatory apparatus, etc.). This fact allow us to compare different ways of “constructing” and patterning organ systems within a set of interrelated animals. We should suggest that, in addition, a significant practical advantage of studying the xenacoelomorphs could be the fact that they seem to possess fewer cell types and organs than most bilaterian animals (lacking proper through-gut, nephridia, complex glands, etc.), making their system more amenable to our future (comprehensive) research efforts. In addition, a better understanding of the genetic control of developmental processes in xenacoelomorphs will provide key insights into the origins and diversification of the bilaterians [10–12].

One of the key innovations linked to the emergence of Bilateria is the origin of centralized nervous systems. How these compact brains are assembled from simpler nerve nets remains a matter of debate that is mostly grounded in the lack of knowledge that we have on the molecular mechanisms that differentially control the development and assembly of nerve nets, cords and compact brains (see, for instance, [8, 12–15]. The comparative approach should provide an answer. In this context the use of members of the Xenacoelomorpha is ideal, since they have different nervous system morphologies (with variable degrees of condensation) all derived from a single, common, ancestor. Briefly, in the group of xenoturbellids, a unique basiepithelial nerve net surrounds the animal body while some acoels have, in addition to a nerve net, an anteriorly concentrated nervous system [1, 16] (for a comprehensive review, see: [17]). The nervous system architecture of the most divergent class of acoels (Crucimusculata) represents, most probably, one of the first instances of the acquisition of a compact brain in bilaterian evolution (nephrozoans would have acquired a compact brain independently). Acoel embryos possess a unique early development program that is known as “duet spiral” cleavage [18]. The first three micromere duets give rise to the ectodermal layer including epidermal and neural progenitors. The formation of the organs’ anlage starts in mid-embryonic stages and the symmetrical brain primordium can be observed at the anterior pole, subepidermally, at early stages [19].

One of the most studied acoel species is *Symsagittifera roscoffensis.* The nervous system of this acoel is arranged in an anterior domain forming a compact brain with neural cell bodies surrounding a neuropil, divided into two lobes and connected by three commissures [17, 20, 21]. The ventral part of the neuropil projects anteriorly to a commissural ring that surrounds the frontal organ. In addition, three pairs of cords arise from the brain and run along the anterior-posterior body axis, in a specific dorso-ventral distribution. Two dorsal cords arise from the posterior part of the brain (specifically from the third commissure), while the remaining four nerve cords lie more ventrally (a ventral central pair and a latero-ventral pair). The most prominent sensory organ is the acoel statocyst, located in the anterior part of the body and surrounded by the brain neuropil. Anterior to the statocyst are a pair of ocelli consisting of several sensory cells and a pigment cell [12, 16, 20, 21]. Juveniles of *S. roscoffensis* are about 220 μm after hatching and the brain occupies more than a third of their body length, a striking difference when compared to the adult specimens, where the brain occupies only a small anterior region (approximately an eighth of the animal’s length) [12, 16, 20–22].

In this context we carried out a systematic characterization of a well-known superfamily of transcriptional regulators, the basic helix-loop-helix (bHLH) proteins. They are widely present in eukaryotes and play an important role in metabolic, physiological and developmental processes [23–26]. Some of them are involved in the development and patterning of the nervous system in many bilaterians (but also in cnidarians (see for instance: [27–29] also reviewed in [26, 30]).

Members of the bHLH superfamily encode for proteins containing a characteristic 60 amino acids long bHLH domain that includes a N-terminal DNA binding basic (b) region followed by two α-helices connected by a loop region (HLH) of variable length. The HLH domain promotes dimerization, allowing the formation of homo- or heterodimeric complexes between different family members. Some bHLHs also include additional domains involved in protein–protein interactions such as the ‘leucine zipper’, PAS (Per–Arnt–Sim) and the ‘orange’ domains [24, 31].

In the past ten years, since the pioneering study of the origin and the diversification of the bHLH carried out by [24], new full sets of bHLH have been identified via the thorough analysis of many metazoan genomes [25, 32–35] (also, in non-metazoans [36]). The complex group of metazoan bHLH transcription factors have been classified, using molecular phylogenetic analysis, into 48 orthologous families (45 different families *sensu Simionato et al.* [25] and three new ones *sensu Gyoja et al* [32]). Based on phylogenetic affinities and general biochemical properties, the orthologous families are organized into six “higher-order” groups named A to F [24, 25, 31, 37, 38]. Group A is especially relevant here since it includes most of the bHLH genes with neurogenic functions in other bilaterians.

Although much has been learnt on the composition and evolutionary pattern of diversification over this last decade, knowledge of patterns of gene expression for individual members (in time and space) remains scant, and mostly focused on a few members of, fundamentally, the A and B superfamilies. In this context, the characterization of full complements of the bHLH and analyses of their expression domains in different metazoan phyla is critical to provide insights understand how these gene superfamilies have changed over evolutionary time. With this general task in mind, our laboratory recently identified the gene members of this group of bHLH genes in two species from the monophyletic group Xenacoelomorpha: the xenoturbellid *Xenoturbella bocki* and the acoel *Symsagittifera roscoffensis* [16] (see Fig. 1 and Fig. 2, for the bHLH members characterized), for which genomes and transcriptomes have recently been sequenced (unpublished).

**Figure 1.**
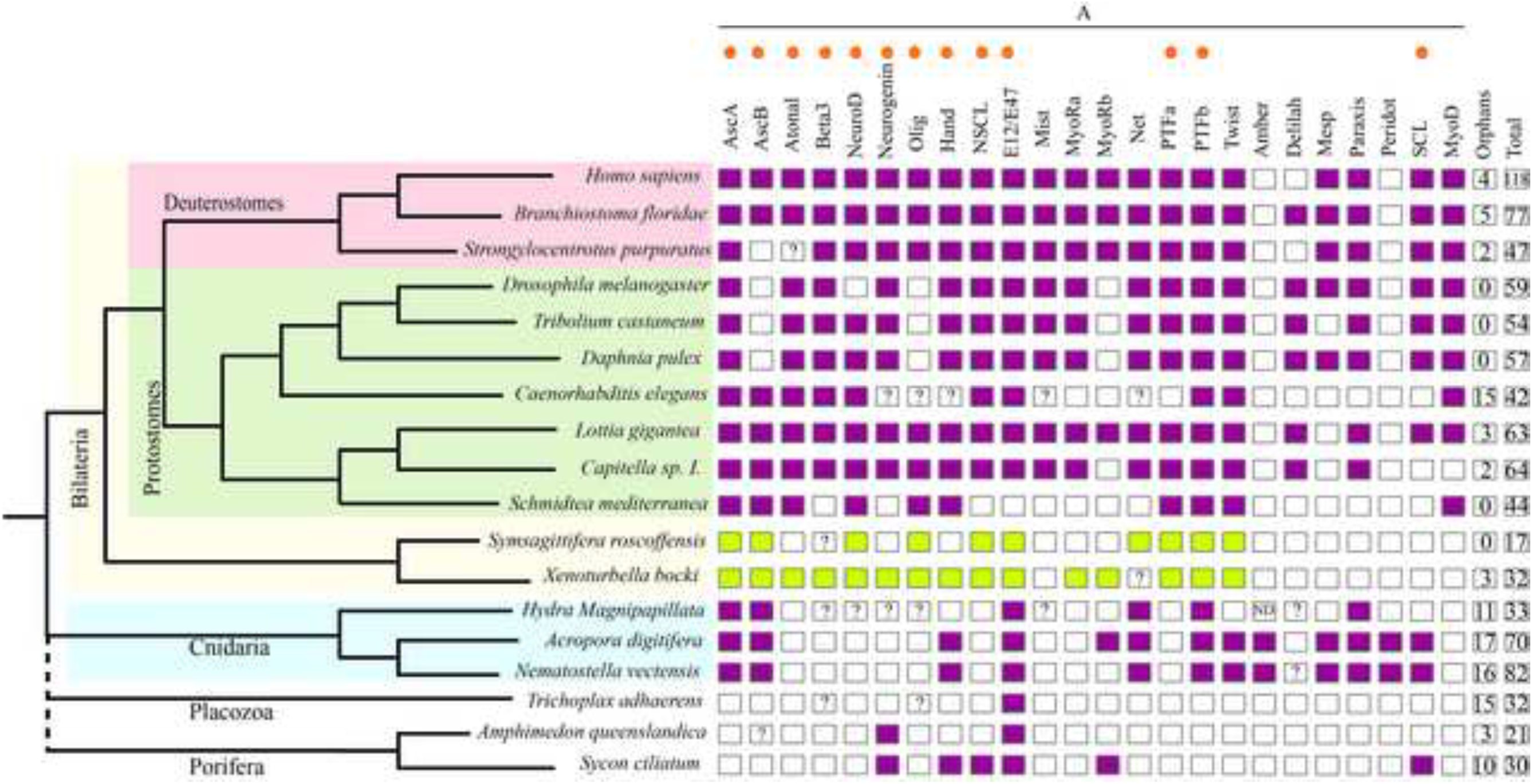
Families from bHLH group A (to which belong most of the so-called ‘neurogenic’ bHLH genes) present in different species from several animal clades. The orthologs’ genes involved in other metazoans neurogenesis and identified in Xenacoelomorpha are indicated with an orange dot. Coloured boxes indicate the presence of the family in that species, while empty boxes indicate their absence. The families from Xenacoelomorpha species are showed with in green. Question marks inside boxes represent the presence of a family member in need of further confirmation (additional gene features). The two last columns represent the number of orphans and the total number of bHLH genes in each selected species (not only from group A). The image allows us to identify the losses produced in the different clades over evolutionary time. The families present in our different species would support the idea of a bHLH gene expansion between the cnidarian and nephrozoan divergence (clearly seen in the group A), as suggested by other authors [25]. Many families have bilaterian, but not cnidarian members; several of them are found in the *X. bocki’s* genome (see also, fig. 2). The data for *X. bocki* and *S. roscoffensis* are derived from our previous analysis [11]. Reference species used here are: *H. sapiens, N. vectensis, Daphnia pulex, Caenorhabditis elegans, Tribolium castaneum, Lottia gigantean, Branchiostoma floridae, Amphimedon queenslandica, Strongylocentrotus purpuratus, Capitella sp. I., D. melanogaster and H. magnipapillata*, all derived from the study of [25]; the data of *Schmidtea mediterranea* was from [33]; the data from *A. digitifera* (plus the latest identifications in *N. vectensis*) and *Trichoplax adhaerens* were obtained from [32, 34]; the *Sycon ciliatum* data was from [35].

**Figure 2.**
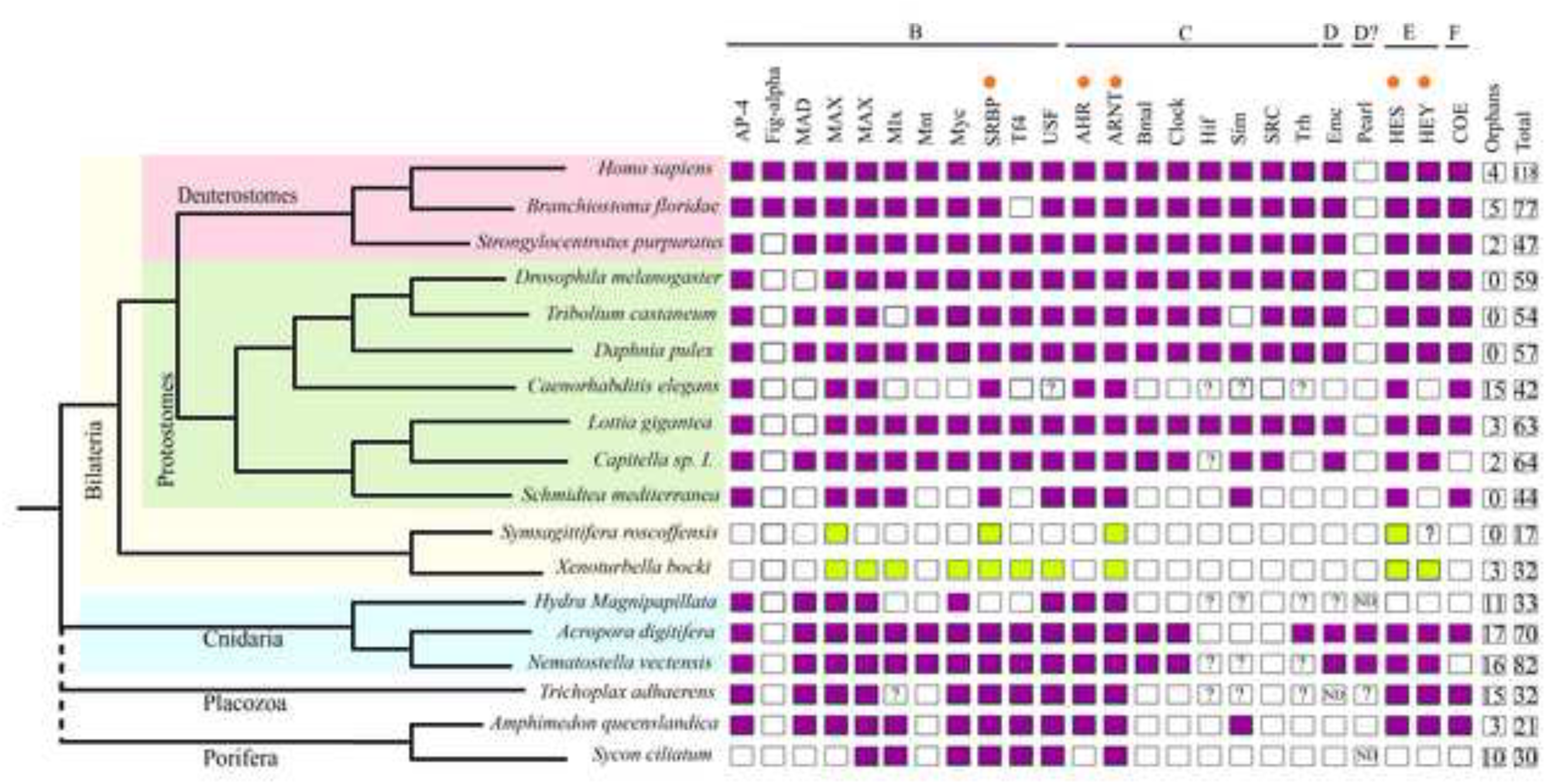
Families from bHLH group B to F present in different species from several animal clades. The gene orthologs with neurogenic role in other metazoan lineages are indicated with an orange dot. Coloured boxes indicate the presence of the family in that species, while empty boxes indicate their absence. The families from Xenacoelomorpha species are showed with in green. Question marks inside boxes represent the presence of a family member in need of further confirmation (additional gene features). The two last columns represent the number of orphans and the total number of bHLH genes in each selected species (from all the groups from A to F). The image allows us to identify the losses produced in the different clades over evolutionary time.. The data for *X. bocki* and *S. roscoffensis* are derived from our previous analysis [11]. Reference species used here are: *H. sapiens, N. vectensis, Daphnia pulex, Caenorhabditis elegans, Tribolium castaneum, Lottia gigantean, Branchiostoma floridae, Amphimedon queenslandica, Strongylocentrotus purpuratus, Capitella sp. I., D. melanogaster and H. magnipapillata*, all derived from the study of [25]; the data of *Schmidtea mediterranea* was from [33]; the data from *A. digitifera* (plus the latest identifications in *N. vectensis*) and *Trichoplax adhaerens* were obtained from [32, 34]; the *Sycon ciliatum* data was from [35].

These genes were cloned and analyzed in the context of a thorough characterization of the bHLH in the xenacoelomorph genomes. In order to characterize the expression patterns of bHLH genes in xenacoelomorphs we started by conducting an analysis of this group of genes in one (experimentally amenable) member of the Acoela, the species *Symsagittifera roscoffensis*, using both coloured *in situ* hybridization (ISH) in embryos and juveniles and double fluorescence *in situ* hybridization (FISH) in juveniles. Our data is also discussed in the context of the evolution of neural system patterning.

## Materials and Methods

### bHLH sequence identification

#### Symsaggitiffera roscoffensis

DNA sequences were extracted from the genome assembly and the embryo (mixed stages) transcriptome and were published in [16]. In the mentioned analysis, sequences were extracted using HMMER (v.3.0) [39] and classified by phylogenetic methods.

### Sampling

Adult specimens of *S. roscoffensis* where collected in Carantec (Brittany, France) during two of their reproductive periods (in April 2016 and 2017). They were cleaned and placed in petri dishes with fresh seawater until they spawned the cocoons in the media. Every cocoon contains several embryos, mostly synchronized (between ten and twenty). We collected the cocoons at different time intervals, with the time when we detected the fresh spawn being the ‘zero’ time. The other times corresponded to the number of hours elapsed between the detection of fresh spawn and fixing the cocoons for in situ analysis. For instance, a sample from 0 to 6 hours contained embryos that developed within the first 6 hours. Embryos were separated from the adults by filtering and then treated using 0.01% Pronase (Sigma) and 0.1% thioglycolate (Sigma) in seawater to permeabilize the egg shells. After cleaning them with seawater, embryos were fixed in 4% formaldehyde (methanol free) overnight at 4°C. A large proportion of cocoons were not fixed, with the aim of obtaining hatchlings later on. These hatchlings were relaxed using 7% magnesium chloride and fixed, as were the embryos. Some specimens were allowed to develop to older juveniles. After fixation all samples were cleaned three times in 1x PBS and dehydrated progressively with a methanol series (25% – 50% – 75% methanol in PBS).

### Gene cloning and in situ hybridization

DIG-labelled and fluorescein-labelled RNA probes were synthesized using the DIG-RNA labelling kit from Sigma, following the manufacturer’s instructions (Id_number11277073910, Sigma-Aldrich). After precipitation, the riboprobes were diluted in hybridization buffer to a final working concentration of 2-1 ng ⁄ μl (depending of the probe). In situ hybridization on whole embryos and juveniles was performed following the protocol published by [21] with a few modifications: 1) the main solvents, PB-Tween and PB-Triton were replaced with TNT (0.1M TRIS-HCl, pH7.5+ 0.15M NaCl+ 0.05% Tween- 20 detergent in RNAse-free water); 2) the proteinase K and glycine steps were suppressed, in order to reduce the damage produced to the samples; 3) we included a step in which we increased the temperature to 80° C during one of the two washes with hybridization buffer (HB), to reduce the background. After this step, we proceed to pre-hybridize overnight. We incubated the sample with the corresponding probe for at least 3 days. The hybridization temperatures were between 55°C and 61°C depending on the probe. The specimens were mounted in 70% glycerol and analyzed using a Zeiss Axiophot microscope (Carl Zeiss MicroImaging GmbH) equipped with a Leica DFC 300FX camera.

Colorimetric *in situ* protocols highlight the domains of highest gene expression. For detailed aspects of the patterns, we rely on the more sensitive fluorescence *in situ* alternatives.

For all clones, sense probes were synthesized that were used as negative controls for hybridization.

### Double fluorescence *in situ* hybridization

Juveniles of *S. roscoffensis* used in FISH analysis needed a photo-bleaching treatment step after the re-hydration. In order to reduce background due to auto-fluorescence, as much as possible, we immersed the specimens in a solution of 1.5% hydrogen peroxide, 5% formamide and 0.5% 0.5xSSC in water (RNAse free), during 15 min and under a white light. Samples were washed twice in PBT (1x PBS+0.1%Triton X-100). The following protocol is a short version of the ISH protocol that it is described in detail in [40]. For FISH all the hybridization probes where diluted to 1 ng/μl. Afterwards, the samples were incubated in anti-DIG-POD 1:500, overnight at 4°C (Sigma-Aldrich, Id_number11207733910). More than four washes, over 2 to 3 hours, in MAB-TritonX-100 0.1%were used to eliminate the rest of the antibody. The signal was developed in TSA red 1:300, in the so-called TSA Buffer (solution of 2M NaCl + 100mM Borate buffer, pH 8.5), over 2 to 6 hours. To stop the development of the signal, samples were washed in PBT. The antibody quenching was made using 1% H_2_O_2_ in PBT 0.1% for 45 min at RT. After washing 2 times with PBT, a 2nd quenching step was done with 2xSSC, 50%Formamide and 0.1%TritonX-100 for 10 min at 56°C. The samples were washed twice in PBT and blocked again, previous to the incubation with the second antibody: anti-DNP-HRP 1:200. After the antibody wash, with MAB-TritonX-100 0.1%, the signal was developed in TSA green 1:300 in TSA Buffer. After the double or single FISH protocols, and in the cases required, we proceeded to combine this procedure with the immunostaining of the samples (using the species-specific anti-synaptotagmin antibody; as a reference, pan-neuronal marker), as explained in the following section.

It is important to note that the general absence of well-defined tissues and the presence of nuclear intermingling in most of the “parenchymal” (external-mesodermal; internal-digestive) tissues of the Acoela makes especially difficult to perform *in situ* hybridization and interpret detailed patterns (as has been noted in other acoel papers).

As in the previous section, for all clones, sense probes were synthesized that were used as controls for hybridization.

### Immunohistochemistry

Immunostaining was performed using the protocols outlined in [1]. *S. roscoffensis* specimens were incubated in primary anti-synaptotagmin (dilution 1:500) antibodies (previously pre-absorbed) and reacted with the secondary antibody [Alexa Fluor goat anti-rabbit 532 (Molecular Probes, Eugene, OR)]. The anti-synaptotagmin antibody was raised in our laboratory using the specific *S. roscoffensis* sequence from a transcriptome analysis (see: [11]). Preimune serum was used as control for all immunochemical experiments.

### Embryo cell counting

Given the difficulty of staging the embryos in the laboratory, we relied, as a good approximation for developmental time, on the number of cells. Embryo cell counting was performed by incubation of all samples for 10 minutes with Dapi. A sample of 12 individuals from a pool of 12 to 24 hours’ post-fertilization embryos were scanned completely using a Leica SP2 confocal laser microscope and their stacks were processed using the software Fiji [41] with the plugin ‘cell counter’ (author: Kurt de Vos; see: https://imagej.net/Cell_Counter). This procedure allowed us to obtain a precise count of the total number of cells per embryo when performing *in situ* procedures.

## Results and Discussion

We carried out a detailed study of the expression of the whole complement of *S. roscoffensis* bHLH genes during development, using early embryos (12 to 24hours post fertilization; embryos have a cells range from 176 to 274, according to our recounts an average of 250 cells/embryo) and hatchlings (time window: 12 to 24 hours post hatch). Taking into account the timing of development of S*. roscoffensis* (and that of other acoels; see [18, 19, 21] for reference time-frames). We selected these stages as relevant starting points for the characterization of nervous system development and hence they should provide us with some initial insights into the roles played by the different bHLH transcription factors in this acoel species. Before discussing further details of the expression patterns, it is first necessary to note here the experimental limitations associated with the acquisition and staging of the embryo samples, which led us to consider time windows of 12 hours instead of exact time points. A few samples were collected within the first few hours after spawning, however the staining patterns for all analyzed genes were either absent or very faint at that stage (irrespective of the probe concentration or staining/developing colour time). It is interesting to note that no expression was visible in many other animal embryos in which bHLH genes were analyzed at the earliest embryonic stages [42–44]. For these reasons we decided not to focus our analysis on earlier embryos (most of the patterns were visible, or more intense, in the biggest embryos at later stages). bHLH expression domains in *S. roscoffensis* were revealed by whole mount colorimetric in situ hybridization (ISH), using probes from 17 different genes (all bHLH genes found in *S. roscoffensis,* as it is represented in Fig. 1 and Fig. 2), in juveniles and embryos. Of these 17 genes, we were not able to obtain expression patterns for ASC_like, NeuroD, PTFa1, PTFa2, PTFb1 or PTFb2 in any of these stages (with the exception of PTFb1 which was expressed in embryos, although the expression levels were very weak). In the case of AscA the signal was only detectable by FISH (not ISH). The absence of expression of PTF family was likely due, to their relatively low expression levels (also correlated with the low numbers of these transcripts in our EST database). We did not succeed with cloning NeuroD likely due to a genome annotation problem. All the other genes, which showed clear *in situ* patterns, are presented in Fig.3. For most of the expressed genes, the detected patterns were always stronger in embryos than in juveniles (presumably also due to the easier accessibility of probes to the interior of the embryos and/or higher relative expression levels). Many of the analyzed genes were expressed in restricted patterns within the embryos, whereas others were expressed more widely in the embryos and/or juveniles. Detailed descriptions of each gene’s expression pattern are given below.

**Figure 3.**
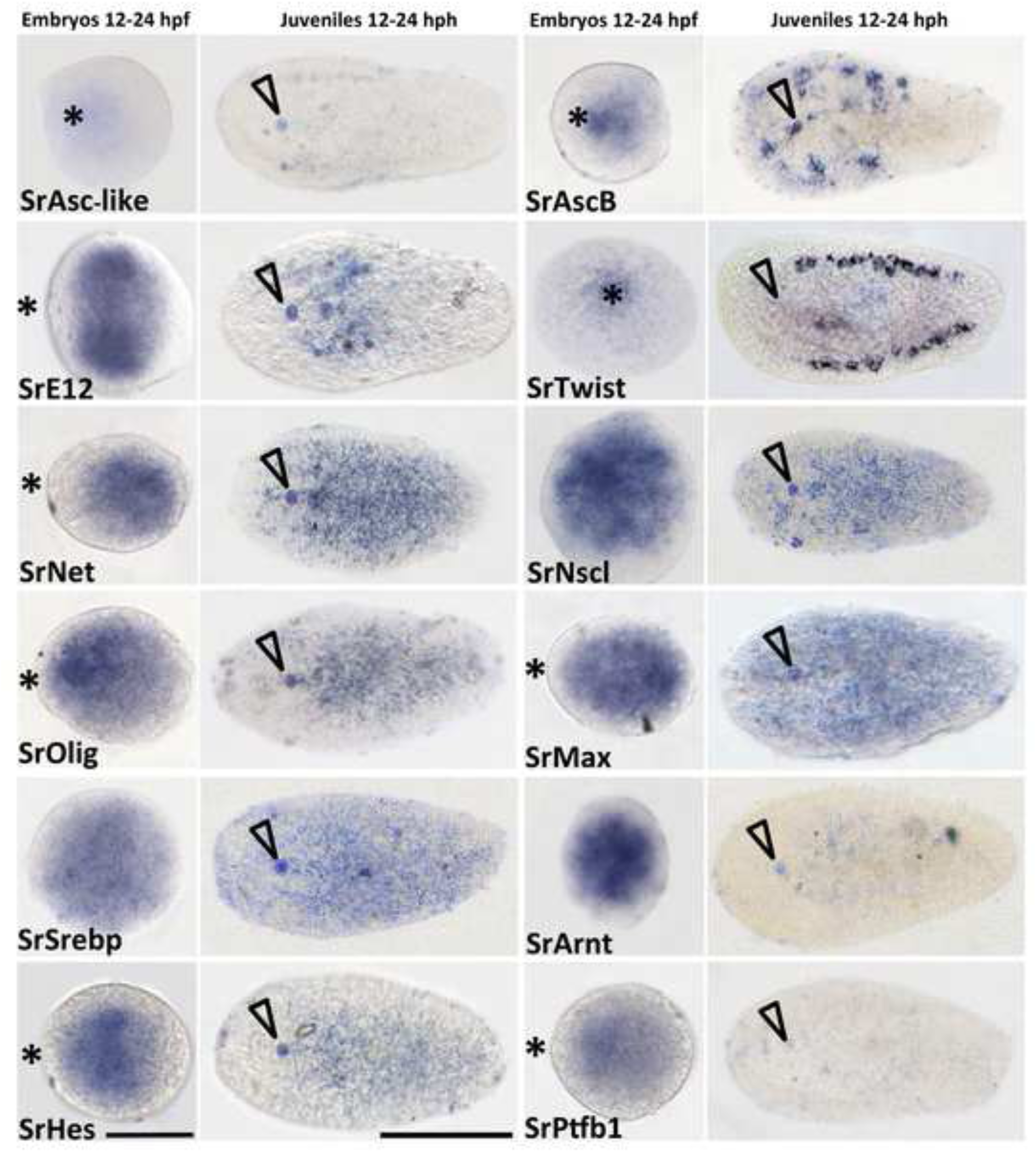
Expression of bHLH orthologs in embryos and juveniles of the acoel *S. roscoffensis*. First and third columns show the expression patterns in embryos, 12 to 24 hours post-fertilization. Second and fourth columns correspond to juveniles from 12 to 24 hours post-hatching juveniles. Names of the corresponding ortholog genes are indicated in each panel. Asterisk indicates the location of the anterior pole of the embryo. Arrowhead indicates the position of the statocyst and therefore the anterior part of the juvenile. Embryos scale bar: 60 um; Juveniles scale bar: 100 um.

It has been known for a while that specific bHLH transcription factors are involved in the process of neurogenesis, especially those belonging to group A. Given the bHLH expression patterns previously obtained by colorimetric ISH experiments, and taking into account their neurogenic role in metazoans, we analyzed some of the bHLH genes in more detail, using double fluorescence in situ hybridization (FISH). This latter approach is more sensitive and allows us a higher resolution analysis of the pattern. The FISH experiments were all carried out in juveniles, with the aim of verifying co-expression domains within the nervous system (we did not succeed with consistent FISH during embryonic stages). To determine the domain of the nervous system, we use a reference marker, the nervous system pan-neuronal gene α-synaptotagmin (see also, in Fig.4, the expression of mRNA and antibody of synaptotagmin are always specifically expressed in the nervous system). However, as α-synaptotagmin mRNA encodes for a synaptic protein (a terminal differentiation marker), we detected no expression in the earliest embryos, prior to 24 hours after fertilization. The first clear signs of the differentiated nervous system were detected, using this marker, in 24–48 hour post-fertilization embryos, showing a bilateral pattern representing the future two anterior brain lobes, the brain primordium (Fig. 4) [12, 16]. In the following paragraphs we discuss the bHLH expression patterns obtained by ISH in the acoel *S. roscoffensis* with a special focus on all genes with a putative function in the nervous system (as reported in other animals).

**Figure 4.**
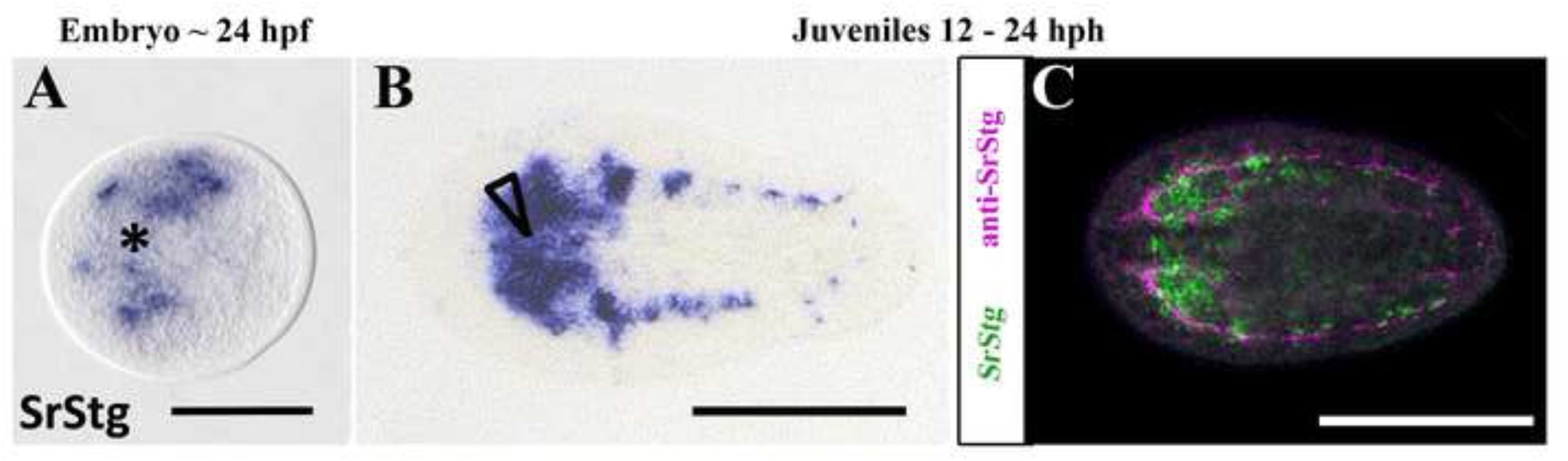
Expression of synaptotagmin gene ortholog in embryo and juvenile of the acoel *S. roscoffensis*. **A.** ISH expression patterns on approximately 24 hours post-fertilization embryos. Asterisk indicates the anterior pole of the embryo. **B.** ISH expression pattern in juveniles from 12 to 24 hours post-hatching. Arrowhead indicates the position of the statocyst and the anterior part of the juvenile. **C.** FISH expression pattern (in green) combined with anti-SrStg antibody (in pink) on a 12 to 24 post-hatching juvenile. Embryo scale bar: 60 um; Juveniles scale bar: 100 um.

### bHLH genes’ developmental expression patterns

Some *S. roscoffensis* bHLH genes seem to share similar expression domains with their homologs in bilaterians, or metazoans, hence pointing to the possibility of a conservation of roles over evolutionary time. In the next section, we analyzed these patterns by following the grouping established for the bHLH genes. We describe here the expression patterns of 13 genes as detected by ISH, with a subsequent, more detailed, focus on those that could be involved in the nervous system development, which were thus analysed by (the more sensitive) FISH Our descriptions are always made in the context of what is known for other metazoan members of the same groups.

### Group A genes families: Achaete-Scute, E12/E47, Twist, Net, Nscl and Olig

Nine genes of the whole *S. roscoffensis* bHLH gene complement were classified as members of the group A: SrAscA, SrAscB, SrAsc_like, SrE12/E47, SrNet, SrNscl, SrOlig/Beta (finally classified as an Olig ortholog) and SrTwist (see [16] for general classification of bHLH genes or Fig. 1 and Fig. 2).

The metazoan Achaete-Scute family transcription factors is divided in two subfamilies: Achaete-Scute A and B. The genes belonging to family A (also named in other clades as “proneural genes”) possess a highly conserved neurogenic role across a wide range of metazoans. It is well known that the Achaete-Scute complex members provide critical proneural function during embryogenesis and the development of adult sense organs in *Drosophila melanogaster* [45]. This role is preserved in other metazoans such as the beetle *Tribolium castaneun,* where TcAsh is also necessary for the formation of the neural precursor [46], and in mice, where the gene Mash-1 is essential for the generation of autonomic and olfactory neurons In the cnidarian *Nematostella vectensis* have shown that the homologous gene NvashA is specifically expressed in a differentiating subset of neural cell types of the embryonic ectoderm [47–49].

When we analysed the spatial domains of expression of the *S. roscoffensis* orthologs of Ash genes we fund two different patterns. The ISH expression pattern of SrAscB was clearly located in the anterior part of the juvenile body and also most likely in the animal pole of the embryo (future anterior part of the juvenile; see also: [19]), We did not detect by ISH the expression of SrAscA (levels too low) in both stages and for the SrASC_like gene (an unclassified member of the Achaete-Scute family see [16]) we detected only a faint expression in embryos. Given the crucial function in the neural differentiation of the SrAscA orthologs (as exemplified in the *Drosophila* case: [50]) and the weak expression of this gene obtained by ISH, we decided to complement our studies by fluorescence *in situ* hybridization (FISH). The mRNA pattern observed by FISH shows expression in an extensive part of the nervous system, revealing the CNS (brain and cords) (Figs. 5A; 5C), the peripheral tracks and the nerve net (Figs. 5B; 5D). The AscA expression domain includes the most posterior part of the cords, the area where they converge (Fig. 5E)

**Figure 5.**
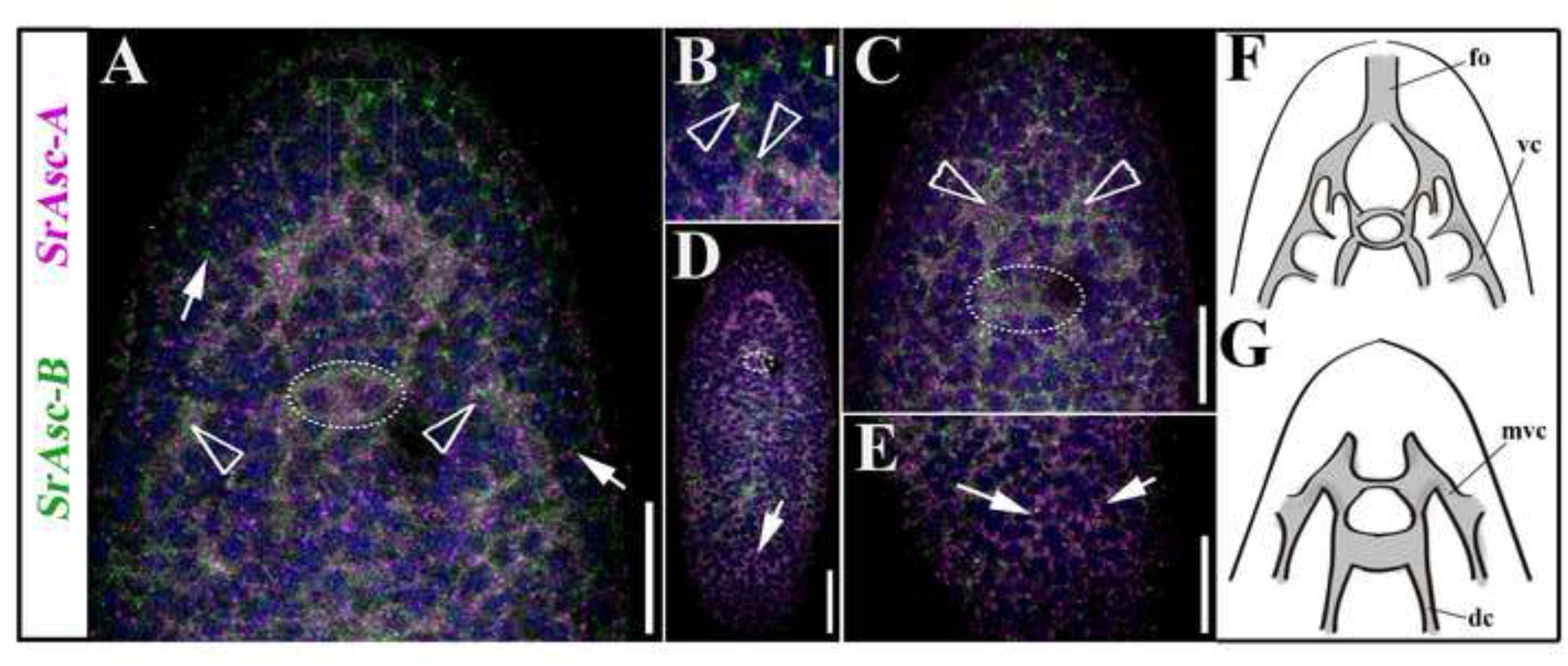
Expression of SrAscA (pink) and SrAscB (green), Achaete-Scute gene family orthologs in (aprox.) 24h juvenile of the acoel *S. roscoffensis*. **A.** Double-FISH expression patterns of a medio-ventral frontal section in the anterior part of the animal. Arrowheads indicate the location of cell clusters where the expression is higher within the CNS (also visible in the SrAscB juvenile’s pattern of the figure 1). Arrows point to the peripheral nerve tracks connected to the animal’s surface. A rectangular box labels the expression domain of both genes in the area of the statocyst. **B.** Detailed image of the double FISH-detected domain of expression in frontal organ’s associated cell populations (arrowheads). **C.** Dorso-medial, frontal, section of the double FISH detecting genes’ expression in the anterior part of the animal. Arrowheads point the location of the two lobes of the brain. The circle surrounds the position of the statocyst. **D**. Dorsal view of a whole mount double-FISH stained animal where it can be appreciated the SrAscA expression in the peripheral nerve net. A circle surrounds the position of the statocyst. The merging of the cords can be detected both anteriorly and posteriorly (arrow). **E**. Detail in a higher magnification image of the posterior part of the organism showed in D, arrows point the posterior end of the cords. F. Scheme of the ventral part of the animal, indicating the nervous system structures where the acoel has the expression domains of studied genes. G. Scheme of the dorsal part of the animal, indicating the nervous system structures where the acoel has the expression domains of studied genes. fo: frontal organ; vc: ventral cord; mvc: medioventral cord; dc: dorsal cord. Scale bars: A=20um; B=5um; C=20um;D=30um.

Different functions have been associated with the Achaete-Scute gene orthologs belonging to subfamily B. These genes play diverse roles and in most cases, they are not directly involved in the development of the nervous system. In fact, most of them regulate the expression of downstream genes in different tissues; for example, the mammalian bHLH Hash-2/Mash-2/Ascl-2 are crucial for development of the placenta [51], the intestinal stem cell fate [52] or the specification of cell types in the immune system [53]. Nevertheless, some patterns seem to be clearly associated with the neural tissue. For instance, the expression of Mash2 (a mammalian ortholog) in Schwann cells or that of the neuronal-specific planarian (*S. polychroa)* ortholog Spol-AscB [54]. In our study, we found that the ISH expression pattern of SrAscB was restricted to a circular area during embryonic development (Fig. 3), while in the juveniles it was localized specifically in the half anterior region of the animal, in two bilaterally symmetric clusters of cells that were connected by a track crossing the anterior-posterior body axis (Fig.3). Analysing this pattern by the more sensitive FISH technique, the cell populations that express SrAscB mRNA clearly form part of the anterior neural cords (Fig.5 and Supplementary material1) though, in addition, they are expressed in some commissures constituting part of the brain. At the posterior end of the expression domain the pattern is shaped as a transversal band of cells, crossing the anterior-posterior body axis at the level of the mouth (also observed by FISH in the supplementary material1). The domains that express SrAscB have a location slightly ventral (supplementary material 1). The nature of these structures (brain, cords and commissures of the nervous system) in the anterior region of the acoel juvenile suggest that SrAscB is expressed specifically in a neuronal population.

Among metazoans, the E proteins play critical roles in cell growth, specification and differentiation, including the neurons. The acoel bHLH Sr_E12/E47 gene showed a high level of expression in two circular domains of the embryo that later on became two anterior-medial regions located immediately posterior to the statocyst (Fig. 3). At first sight, our results seem to differ from those obtained for the E12/E47 gene orthologs in, for instance, *H. sapiens* and *Drosophila* (Daughterless), in which expression is found in most tissues. However, we should point out that different studies have found an increment in the mRNA expression of E12/E47 (and other E2A mRNAs) in some areas of rapid cell proliferation and differentiation in several tissues, including neural tissue. These levels decrease progressively during neurogenesis, and become almost undetectable in the adult nervous system [55]. This is consistent with our observations, with the strongest expression pattern in embryos at around 24 hpf, and lower levels in the anterior body of the juvenile (Fig. 3). The expression domain in juveniles is in an area partially overlapping the location of the brain (Fig. 3 ISH). The pattern is consistent with an early role in the development of neurons described for other organisms [56], putatively in populations of non-terminally differentiated neurons or neuronal progenitors. E12/E47 genes form heterodimers with other group A bHLH factors [24, 57]. In fact, some authors have suggested a specific role of E-proteins in early neural differentiation [58] and, in the same vein, recent studies have confirmed that E-proteins orchestrate neural stem cell lineage progression [56]. bHLH genes encoding other E proteins with a similar role are seen in, for instance, the planarians, in which the gene e22/23 is expressed in the CNS [33]. What seems clear is that both genes Sr_AscB and Sr_E12/E17 are most probably associated with the development of the nervous system.

In contrast to the mRNA expression patterns found in the above members of group A, the other members of this group (SrTwist, SrNet, SrNscl and SrOlig) showed diverse expression patterns mostly in the middle body region. SrTwist was expressed in a discrete spatial domain well delimited in the embryo’s animal pole and organized as a pair of bilateral bands on both sides of juvenile specimens. This finding is in agreement with a previous studies from our laboratory that analyzed bilaterian mesodermal gene expression have already revealed the expression pattern of twist orthologs in embryo, juvenile and adult stages of *S. roscoffensis* [59] and adults of the acoel *Isodiametra pulchra* [60], suggesting its expression in part of the gonads, the male copulatory organ (only in *I. pulchra*) and neoblasts, all of which are mesodermal derivatives. Furthermore, the mesodermal role of twist homologs, for instance, in *Drosophila*, is well known [61].

The expression patterns of the remaining three genes, SrNet, SrNscl and SrOlig, showed a scattered distribution throughout the body though slightly different from each other. The expression domain of SrNet gene is a bit larger than that of SrOlig and SrNscl (Fig. 3). In the case of *S. roscoffensis* SrNet was expressed in a region located in the middle of the embryo. In juveniles, the SrNet expression signal covered the statocyst, forming different lines in the anterior territory of the body, with a high expression in the posterior half of the body (Fig. 3). In agreement with this SrNet expression pattern, and taking into account the roles described for Net homologs in other animal models (for instance in the jellyfish *Podocoryne carnea*, Net is expressed in the entocodon, a mesoderm-like structure that gives rise to the striated and smooth muscle of the bell [62] and in *Drosophila* DmNet is required to maintain the inter-vein regions during development [63]) we suggest that the acoel Net homolog might have a mesodermal role, without any significant participation in neurogenesis, although clearly further studies are required.

Expression of SrOlig in 12–24 hour post-fertilization embryos of the acoel occurred in a scattered pattern covering most of the embryo, however we (qualitatively) detected the highest level of expression in a smaller localized region (Fig. 3). Expression was mostly limited to the center of the juvenile body, being very low in the zone of the brain and with no signal in the posterior part of the organism (Fig. 3). FISH experiments show low expression in the brain and a zone corresponding to the region where the brain cords converge anteriorly (Fig. 6A, 6B). Additional to the central expression found in juveniles, SrOlig seems to be more expressed dorsally and most probably present in the nerve net and in a low range in the cords (Fig. 6A, 6B). Due to the conserved role of this gene family among different clades (mammalian factors Olig1 and Olig2 are involved in the specification of progenitor populations that produce motor neurons and later oligodendrocytes (reviewed in [30, 64–68])., and taking into account the absence of expression in the region anterior to the statocyst, where most of the nervous system of the acoel is located, we should be cautious about stating the specific cell populations that express SrOlig. Noteworthy is that similarities were found between the pattern in *S. roscoffensis* and that in the planarian *S.mediterranea*; in both cases a small anterior population of cells, separated from the rest of the domain, expressed the respective Olig orthologs.

**Figure 6.**
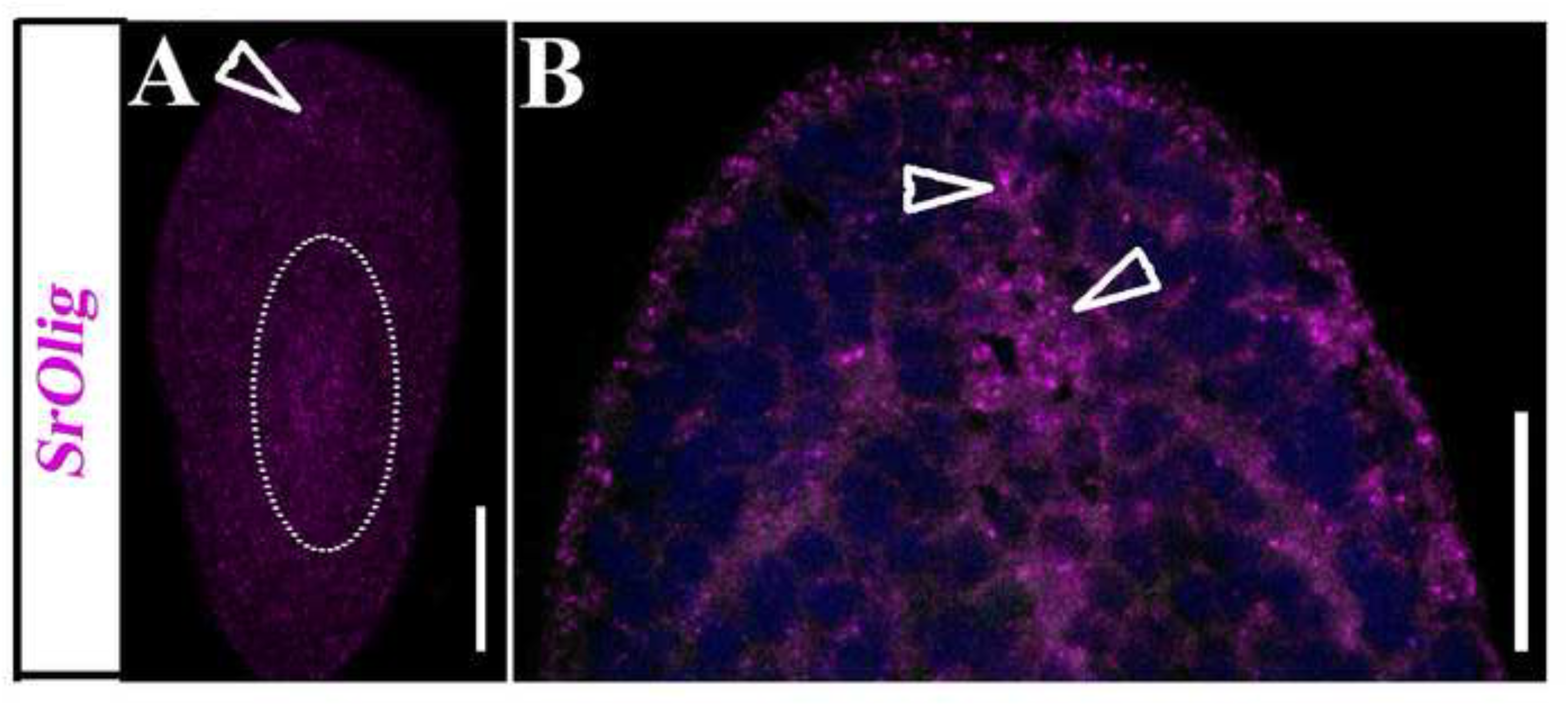
Expression of SrOlig (pink) gene, in approx. 24h juveniles of the acoel *S. roscoffensis*. **A.** Dorso-frontal section of a whole mount single FISH stained animal showing the SrOlig gene expression, in a small, concentrated, domain located in the frontal area of the brain (arrowhead) as well as in the area near the mouth (circle). B. Ventral section of a whole-mount double-FISH stained animal showing the expression pattern of SrOlig at a higher magnification; arrowheads point to the expression in the brain area. Scale bars: 40um.

Among metazoans, the contribution of Nscl gene orthologs to the development of sensory cells and neurons has been well-described [69, 70]. The *S. roscoffensis* bHLH gene SrNscl showed, in the juvenile, a very low expression, in a domain that seemed to be entirely posterior to the statocyst (Fig. 3). Interestingly this gene showed, however, what seemed to be a strong expression in an extensive region in our embryos (compared with the expression levels, at similar times and probe concentrations, of other bHLH). Nscl mouse’s orthologs show strong expression during the post gastrulation period, most likely due to their role in early neurogenesis [71]. Functional analysis of the Nscl-1 mouse ortholog demonstrated the need for this gene for correct neural cell differentiation, and in combination with Nscl-2, it is required to control the migration of neuronal precursor cells [72, 73]. With the data obtained, and due to the low expression levels in juveniles, we cannot conclude with certainty that the expression of SrNscl occurs within the nervous system.

### Group B gene families: Max and Srebp

The bHLHs genes belonging to group B in *S. roscoffensis* are SrMax and SrSrebp. The metazoan orthologs of Max are involved in cellular proliferation, development and differentiation. Several studies have shown that the MAX protein forms heterodimers with other transcription factors such as the bHLHs MYC and MAD [74]. MYC-MAX heterodimers are involved in the transcriptional activation of different target genes [75, 76]. In *S. roscoffensis*, MAX is the only known member of these complexes, since MYC, MAD and other related bHLH transcription factors such as MLX were not found in the genomes or transcriptomes [16]. This is interesting in itself since it suggests a developmental function for Max that is independent of heterodimerization. Expression of SrMax mRNA in juveniles is completely ubiquitous; however, this was not the case in embryos, where, even though the signal covered a large part of the embryo an enriched expression domain was still observed (Fig. 3).

The other member of group B in our species is SREBP. Srebp orthologs are involved in animal homeostasis, in the regulation of sterol metabolism. They regulate the gene expression of most of the enzymes involved in cholesterol biosynthesis. SrSrebp expression in *S. roscoffensis* is widespread, at all stages analyzed (Fig. 3) consistent with having a similar (generalized) role in metabolic regulation, perhaps including supporting glial cells (see [20] for evidence of glial cells in *S. roscoffensis*).

In the case of mice and rats, the Srebp gene orthologs are also expressed in several cell types, including astrocytes, oligodendrocytes and Schwann cells, those that are very active in lipid metabolism [77]. Moreover, in the adult planarian *S. mediterranea* this gene shows intense expression over the whole body, where it is expressed as well in different cell types [33].

### Group C gene families: ARNT

The only representative of group C in our acoel species is ARNT (aryl hydrocarbon receptor nuclear translocator). ARNT can form heterodimers with several bHLH proteins and its function depends on its dimerization partners. For this particular reason, members of the ARNT family tend to be widely expressed [78]. Specifically, the AHR/ARNT system controls processes such as oxidation/anti-oxidation, epidermal barrier function, photo-induced response, melanogenesis, and innate immunity [33, 42] [79]. In our genomic/transcriptomic analysis, we have found only the gene encoding for ARNT transcription factor, and not the gene for its partner AHR, contrary to what has been described in most metazoans [16][3, 5]. SrArnt mRNA was expressed in the entire embryo with small areas inside the domain showing higher levels of expression (Fig. 3). In juveniles, probably due to the low level of signal, we detected a very faint expression, widespread in the body (Fig. 3), suggesting that ARNT in *S. roscoffensis* is probably expressed in a variety of cell populations.

### Group E gene families: HES/HEY

As for the previous class, *S. roscoffensis* group E comprises a single HES/HEY subfamily member. The SrHes/Hey gene was expressed in the anterior-medial region of the embryo and later on, in the juvenile stage, its expression is lower with an area a little bit more intense in the central part of the body, posterior to the statocyst (Fig. 3; Supplementary material 2). A detailed expression analysis, using FISH, locates the main domain of SrHes/Hey expression dorsally, in the cords (supplementary material 2; Fig. 7A, 7C, 7D) and mid-ventrally, at a lower level, within the brain (Fig. 7B, 7E).It is well known that HES proteins act by inhibiting proneural bHLH protein functions through a mechanism that involves the repression of proneural gene expression (reviewed in [80]. A reduction in the number of neural progenitor cells occurs in the absence of mouse Hes genes, in parallel with the premature neural differentiation of neuroblasts [81, 82]. Moreover, knockdown of its planarian ortholog hesl-3 during regeneration leads to a reduction in the neural population and a miss-patterned brain [33]. The expression pattern obtained for the SrHes/Hey gene in juveniles of *S. roscoffensis* resembles that of the three Hes gene orthologs in planarians (hes-1, hes-2 and hes-3), which are not expressed in the most anterior part of the body but are otherwise highly expressed in the central part of it, although only hes-3 is clearly expressed within CNS [33]. In spite of the specificities of each ortholog in each biological system, the expression of Hes genes in neural progenitor cells seems to be well conserved among metazoans, and this role is compatible with our data.

**Figure 7.**
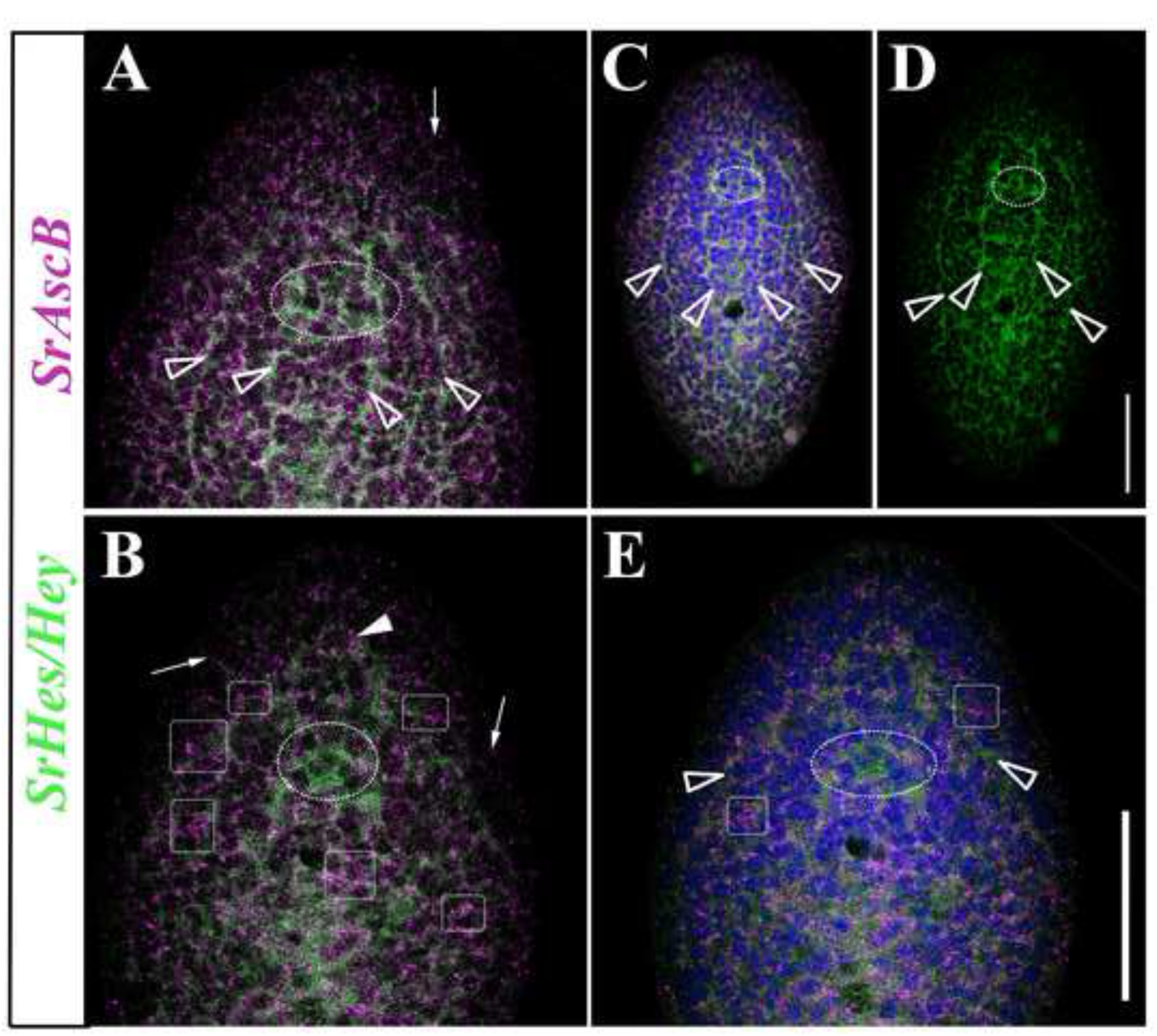
Expression domain of SrAscB (pink) and SrHes/Hey (green) genes in, approx., 24h juveniles of the acoel *S. roscoffensis* (several panels). **A.** Dorso-frontal detail of the anterior part of the organism where it is detected the strongest expression of AscB (pink) in the nerve net and some peripheral tracks (arrow). SrHes/Hey shows expression in the dorsal and medio-ventral nerve cords entering the brain (green) (arrowheads). Also some commisures connecting the brain can be here appreciated. **B.** Dorso-frontal detail of the anterior part of the organism where it is detected the expression of AscB (pink) in the nerve net and in the cross commissures (squares) plus in some peripheral tracks (arrows). SrHes/Hey shows expression in the origin of the dorsal cords, near to the statocyst, and in the ventral nerve cords entering the brain (green) until connectUntil connect what? (arrowhead). Also some commisures connecting the brain are detected. C. Dorso-frontal view of a whole specimen where it is detected the expression of AscB (pink) and SrHes/Hey (green) with DAPI. Arrowheads point the expression of SrHes/Hey in the cords. D. SrHes/Hey (green) expression with arrowheads pointing the neural cords. E. Same figure of panel A with DAPI, arrowheads pointing the ventral nerve cords. A circle surrounds the statocyst in all the panels. Scale bars:40um

### Summarizing the Developmental expression of neural bHLH transcription factors in ***S. roscoffensis***

As stated in the introduction, several studies in many animal systems have demonstrated the involvement of different bHLH family members in neurogenic processes, with the characteristic that most of them (but not all) belong to the so-called group A. *Symsagittifera roscoffensis* possesses an interesting set of bHLH genes, some of which are expressed in domains clearly overlapping the anterior part of the nervous system: SrAscA, SrAscB, SrE12/E47, SrHes/Hey and SrOlig. Their relative expression domains are represented in a schematic model of the juvenile acoel (see Fig. 8). The differential patterns of expression of these bHLH genes in different parts of the nervous system suggest the diverse roles that these genes may have in the patterning of the nervous system and the development of its final architecture. These findings are consistent with those of other functional studies, which point to the importance of the combined expression of some bHLH transcription factors in patterning the neural tissues. They act in concert (or downstream) with other patterning genes that provide positional identity along the major body axis of animals, for instance in the dorso-ventral axis (Pax, Nkx and Irx) and along the anterior-posterior body axis (Otx, Gbx, En, and Hox families) (reviewed in [80]; [15]; our unpublished acoel results). Moreover, some studies have reported the combinatorial activity of orthologs of bHLH proneural proteins (Mash1, Hes1, Olig2) with other patterning proteins such as Pax6 and Nkx2.2 promoting cell type specification in mice (see for instance: [68]).

**Figure 8.**
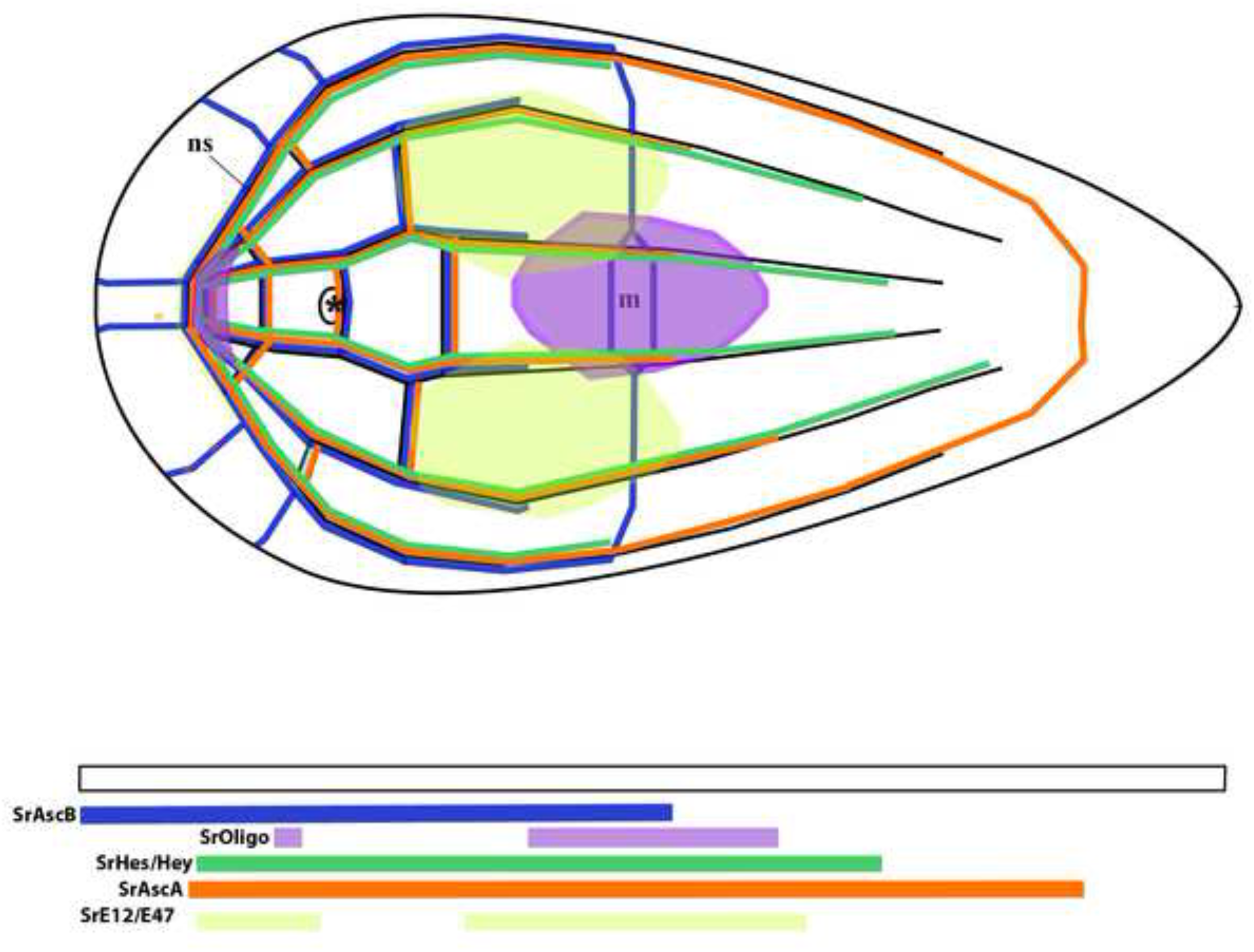
Schematic model of all collected mRNA expression patterns obtained in our study of the bHLH genes (with a focus on the expression domains within the nervous system). Abbreviations: **ns:** nervous system; **m:** mouth; **(*):** anterior statocyst. The model is based, essentially, on all the FISH (single and double) *in situ* data, since they provide us with higher, more accurate, information on relative spatial patterns.

In the acoel nervous system a similar system of neural specification seems to be in place, where a subset of the bHLH genes are used in different (and overlapping) neural domains. We are aware, though, that this is a preliminary characterization of the whole superfamily of genes and that assigning specific functions to genes or combinations of them is still premature, in absence of more detailed and /or complementary data, such as Q-PCR and in situ hybridizations on histological sections. However, this study provides us with the cartography of neural expression domains, a roadmap to further investigations.

### A final caveat: The bHLH complement in the Acoela

The complement of bHLH detected in the acoel *S. roscoffensis* is reduced in comparison with other metazoan organisms studied. Most of the described roles of the bHLH orthologs in other organisms could not be performed with the small complement of bHLH present in *S. roscoffensis*. There are several different possible reasons for the detection of a reduced set of bHLHs in this clade. First, there is the obvious possibility that some bHLH appear to be missing because their sequences are too divergent and therefore, difficult to identify (see Fig. 1 and Fig. 2, for a schematic representation through the different phyla). This is not unlikely in acoels, as their genomes have clearly changed a lot because of the high rate of sequence evolution. This has generated some clear genomic modifications, among which genetic losses are common. A reduction in the number of protein complements has been reported in several families previously [12, 16]. Other possible factors are the highly divergent sequences of members of other families, for instance the sequences belonging to the Wnt family of ligands identified in *S. roscoffensis* and the acoel *Hofstenia miamia*. They are clearly derived and this has made it impossible to classify them into the well-known metazoan families [12, 83]. An alternative possibility for the scarce number of bHLH relatives in the acoel genome is that we are using a newly sequenced (draft) genome, which we assume is almost complete, but could still be missing some fragments of the genome. In this context we should mention that the parallel use of transcriptomes, from adults and embryos, has not provided any new sequences that were not present in the genome.

## Conclusions

The genome of *S. roscoffensis* contains a relatively low number of bHLH transcription factors. However a careful phylogenetic analysis of the sequences has revealed that this clade has bHLHs from various subfamilies. Acoels possess, at least, a bHLH belonging to five of the six classical high order groups. The conservation of the bHLH sequences analyzed is also paralleled by the conservation of the different gene expression domains. Clear correlations can be made between the expression domains in acoels and those reported for other metazoans, suggesting conservation of roles over evolutionary time. Our study detected the presence of a pool of bHLH genes (SrAscA, SrAscB, SrE12/E47 SrOlig and SrHes/Hey) with expression patterns specific to territories within the nervous system, most probably in different cell populations of neurons and/or neural precursors. Their expression, which starts in most cases during embryogenesis, suggests that they are involved in the early specification of neural precursors and the later formation of the nervous system (or subdomains thereof). This analysis, together with that of other genes studied previously in our laboratory, such as the Hox family members Cdx and SoxB, constitutes a first step towards a clear understanding of how the nervous system is assembled in acoels. The study of many other regulatory genes in our laboratory (unpublished) suggests a complex gene network controlling the development of the acoel nervous system. The further characterization and detailed knowledge of the genes involved in neural development and patterning will help to address specific developmental issues in the future; issues that would be impossible to tackle in absence of this cartography of expression domains. Having access to the sequence of the *S. roscoffensis* genome has been the key factor in this study.

## ACKNOWLEDGEMENTS

We would like to acknowledge the support offered by Drs. Cristian Cañestro (UB; Barcelona) and Anna Aragay (IBMB-CSIC; Barcelona), which was critical in order to finish this research project. E. Perea has been a recipient of an APIF (UB) doctoral fellowship. We should also acknowledge the continuous help provided by Brenda Gavilán, Beatriz Albuixech and Carlos Herrera during most of the duration of this project. We greatly appreciate the permanent support of the members of the CLSM facility of the University of Barcelona.

PM would like to acknowledge Prof. William McGinnis (UCSD), in which laboratory he learned the use of basic tools for multiplex in situ hybridization.

Sequences and alignments are publically available through the NCBI. The authors declare that none of them have any competing interests.

We would like to thank the referees for their thorough, constructive, comments to this paper. They have helped us improving the manuscript.

## Supplementary material

**Supplementary material 1: Detailed expression of SrAscB (pink), Achaete-Scute gene family ortholog B in (aprox.) 24h juvenile of the acoel *S. roscoffensis*.** Panels correspond to three different planes along the dorso-ventral axis. Scale bar: 20 um.

**Supplementary material 2: Detailed expression domains of SrAscB (pink) and SrHes/Hey (green) genes in, approx., 24h juveniles of the acoel *S. roscoffensis*.** The different rows correspond to their different planes along the dorso-ventral axis of the juvenile. In the bottom line of every column it is indicated the gene combination used, except in C, C’ and C’’, which correspond to Dapi stainings. Arrowheads point to the expression domain within the nervous system. Scale bar: 40 um.

