## Supplementary figures and images for "Characterization of the bHLH Family of Transcriptional Regulators in the ACOEL *S. roscoffensis* and their Putative Role in Neurogenesis"

### Supplementary Materials

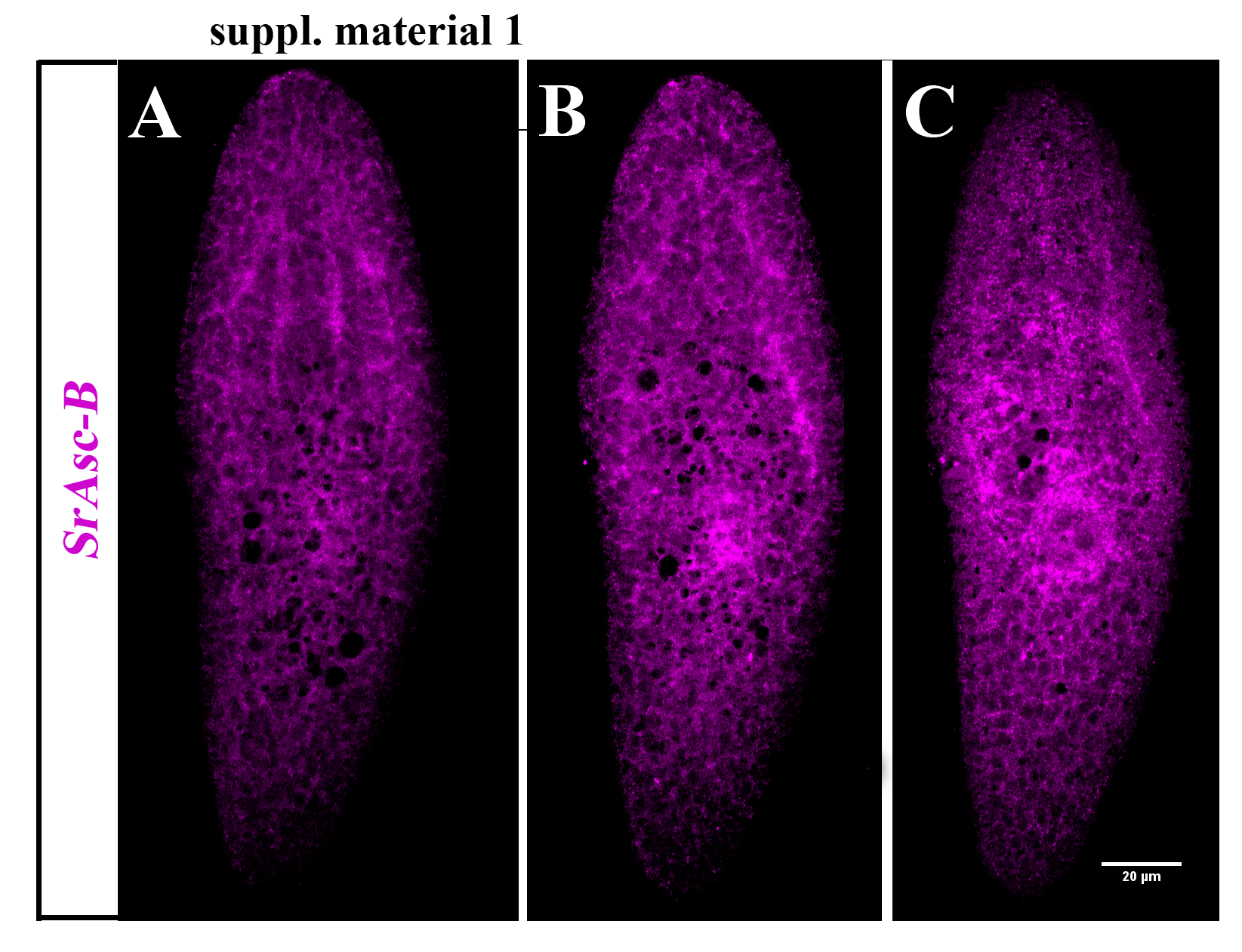
